# Caste-specific processing of construction pheromones in mound-building termites

**DOI:** 10.1101/2025.09.08.674840

**Authors:** Sree Subha Ramaswamy, Sanjay P Sane

**Author notes:** **Competing Interest Statement:** The authors declare no conflict of interest.

## Abstract

Mound-building termites construct massive structures from soil. These structures contain intricate, interconnected passageways linking multiple chambers and atria, which house the brood and fungus gardens that support wood digestion, while also enabling efficient movement of termites. Remarkably, the major and minor workers who build these structures in their dark, subterranean milieu lack image-forming eyes. Coordinated construction on such a scale requires close communication among thousands of individuals. How do termites communicate and coordinate their activity within the mound? Previous studies have shown that communication between termites is mediated by a range of pheromones. A prominent hypothesis, the *stigmergy hypothesis,* suggests that termites communicate with their nestmates indirectly, by embedding chemical signals in the soil. We developed a *choice assay* to measure termite attraction to soil containing termite-deposited cues, offering a choice between soil from a repair site and unprocessed control soil. Using physical and chemical extraction methods, we removed volatile and non-volatile components and then measured termite attraction to the treated soils. In these experiments, termites preferred soil from their own mound’s repair site over surrounding soil, indicating the presence of embedded pheromones. Of these, the non-volatile components could elicit responses to soil preserved for long periods of time. When extracted and added exogenously, these pheromones made unprocessed soil attractive to termites. Major and minor workers responded to volatile and non-volatile cues differently, indicating caste-specific responses to pheromones. These data show that termites use soil as a medium for communication, consistent with the *stigmergy hypothesis*.

**Significance:** Mound-building termites achieve the collective construction of elaborate structures through finely tuned coordination of their building behaviors. Lacking image-forming eyes, they rely heavily on a sophisticated chemical communication system by embedding pheromones in the soil that their nestmates sense and interpret, to coordinate behavior. To investigate this communication system, we conducted a series of experiments using a novel *choice assay* which measures the behavioural responses of termites to pheromone cues in the soil. These pheromones contain volatile and non-volatile components to which the termites respond in a caste-specific manner. The non-volatile components elicit responses over a very long period. Our findings provide new insights into the chemical communication that drives collective behavior in social termites, with implications for collective robotics.

## Introduction

Social insects including bees, ants, and termites employ complex systems of communication to coordinate cooperative building, navigation, and environmental interactions, all processes vital for maintaining large colonies (1–4). Unlike most social insects, which are Hymenopterans, termites form a monophyletic clade within the cockroach order Blattodea (5). Among them, mound-building termites construct extensive structures critical for thermoregulation, ventilation, fungal farming, and humidity regulation within their nests (6–11). Their mounds house entire colonies of termites across generations and have been likened to large human-built structures, which must also resolve similar challenges. Unlike human-built structures, however, termite mounds are constructed in a decentralised manner, relying on local communication to coordinate their building activities. Although mound-building termites lack image-forming eyes, they communicate efficiently with their nestmates using chemical and acoustic cues(12, 13). This enables them to recruit nestmates to specific sites across extended spatial and temporal scales, and cooperatively build, repair or maintain their nests. The pheromone-mediated communication system of termites is thus an excellent system in which to study how social insects generate, encode and decode these chemical signals, which is a central question in sociobiology.

In his classic paper, Grasse proposed the *stigmergy hypothesis,* which argued that social insects including termites, communicated with each other *via* signals embedded in the structures that they built (14). The chemical cues underlying mound construction likely involve multiple pheromones and receptors that elicit role-specific responses in termites. These cues are hypothesised to elicit local actions, such as trail-following, nest-building, defence, and reproduction (13, 15–18), giving rise to large, emergent structures (19, 20). Although the *stigmergy hypothesis* offers a conceptual framework to understand termite communication, its underlying mechanisms remain largely unknown.

To address these questions, we studied the building behaviour of the mound-building termite *Odontotermes obesus* (Movie S1), which occurs commonly across the Indian peninsula.

These mounds can rise to a few metres above ground with comparable depth underground (Fig 1A). In *O. obesus,* mound-building is carried out by major and minor workers, who are distinct in their morphology (Fig 1B). They are protected by soldiers with large mandibles adapted for defence. The mounds are lined with complex passageways and chambers housing the queen, king, their brood, and a special *Termitomyces* fungus garden that helps the colony digest wood(21).

**Figure 1:**
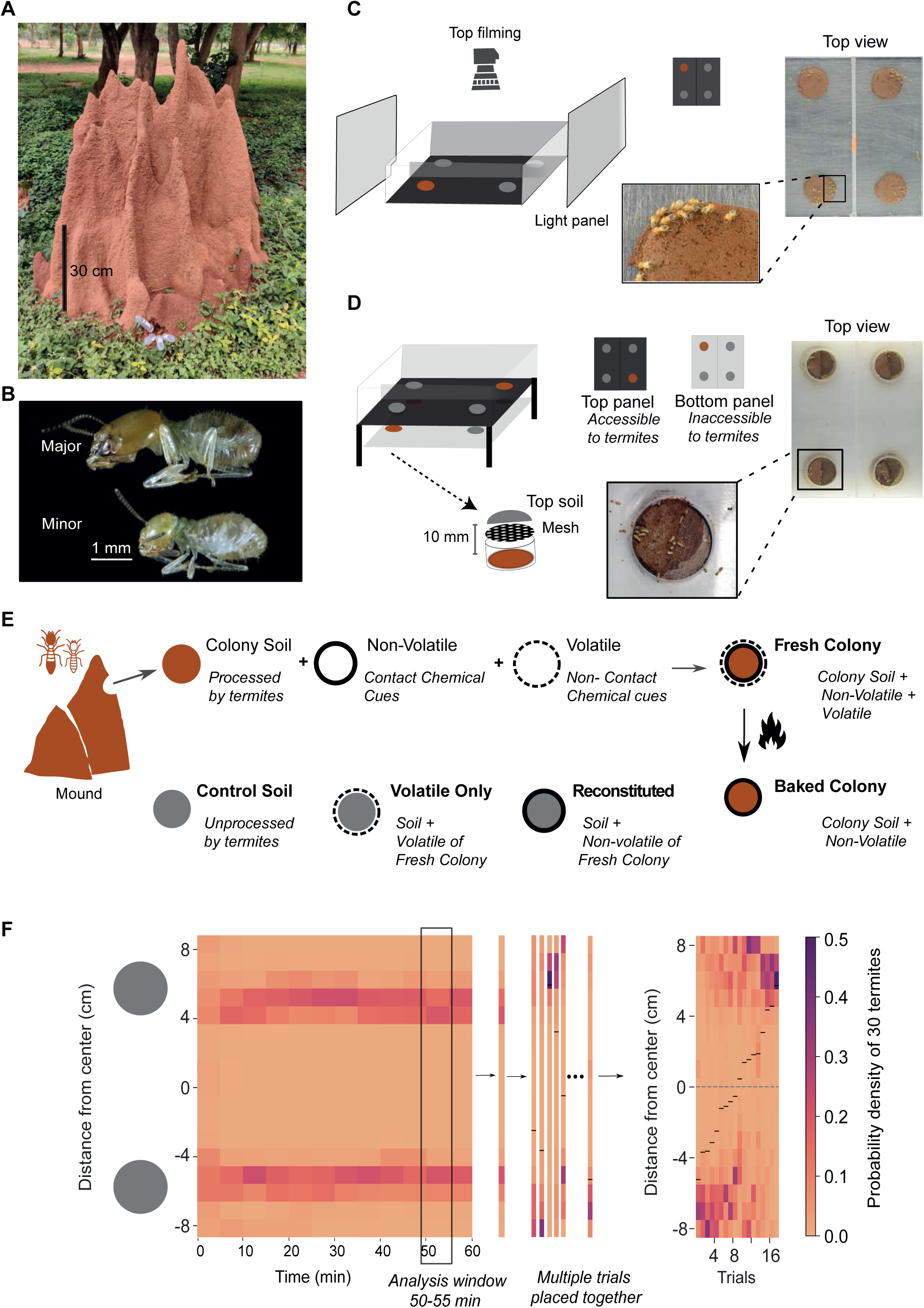
Assay design and visualisation framework for caste-specific termite responses to soil cues. **A)** An *Odontotermes obesus* mound with a visible injury at the base, facilitating access for termite collection. **B)** Representative images of a major (top) and minor (bottom) worker termite, illustrating caste differences. **C)** Schematic representation of a choice assay experiment with two wells. The left well has a treatment patch on the top and a control patch on the bottom. The right well contains two identical control patches to test for side bias. The arena is illuminated from both sides and filmed from above. Termites are introduced at the centre of the arena. The top view (at steady state, 55 min) shows termite distribution over soil patches. A close-up highlights termite aggregation on a patch. **D)** Schematic of a mesh choice assay. Two semi-circular soil patches are placed on the top panel, separated by a mesh that prevents direct contact with soil beneath. Test soil is placed in a bottom panel, ∼10 mm below the mesh, to deliver only volatile cues. Termites are introduced at the centre on the top panel. The top-view image shows termite distributions at steady state, with a close-up of the mesh structure. **E)** Iconography key used across schematics and plots. **F)** (Left to right): The arena width is collapsed, retaining only the Y-axis, which is binned into 1 cm segments. The resulting heatmap represents termite presence along the Y-axis over time (X-axis), with each vertical strip representing a time slice. The colour bar indicates the probability density of termites at different Y-positions. Grey shaded regions represent the locations of the soil patches on the Y-axis: from 4 to 8 cm and from -4 to -8 cm. The 50th-55th minute time window is used for steady-state analysis, and median positions for each trial are marked with a black line. For multi-trial visualization, multiple such strips (one per trial) are stacked and sorted by ascending median, forming a tapestry plot.

To study their communication during building, we developed a novel lab-based assay which measured the attraction of termites to soils that were previously processed by their nestmates, and infused with pheromone cues (Fig 1 C, D). In this *choice assay*, termites were offered a pair of soil patches made out of various soils that were either unprocessed, termite-processed or heat- or chemically-treated (Fig 1E, Table S1), and their relative attraction to these patches was recorded using an overhead camera which tracked the 2D movements of the termites within the arena (Fig 1C). Their movements were then plotted as heatmaps that enabled quantification of their choice (Fig 1F). This assay demonstrated that termites use specific pheromone cues to stimulate building activity in their nestmates. These cues contain volatile and non-volatile components which generate caste-specific responses in major and minor workers.

## Results

### Termites are attracted to soil from the site of repair

Using the *choice assay*, we tested the hypothesis that termites embed chemical or textural cues in soil during construction to attract and recruit nestmates to the repair site. If true, more termites should be attracted to a patch of freshly processed soil from a repair site (*Fresh Colony soil*, see methods) as compared to *unprocessed (Control) soil*. In contrast, they should show no preference between the two control soil patches. Our observations were consistent with these expectations (Fig 2). In experiments with two identical control soil patches, major (Fig 2Ai) and minor (Fig 2Bi) workers were not preferentially attracted to either patch (p=0.82 for majors, p=0.22 for minors, Wilcoxon signed-rank test, Table S2) and distributed themselves across both patches (39 trials for majors with 1170 termites, and 17 trials for minors with 510 termites; see Fig 2Ai, 2Bi, 2C, Table S2 and Table S3 for statistical analysis). In contrast to the control soil patches, they were attracted to the *Fresh Colony soil* (Movie S2) over *Control* soil in both major (20 trials, 600 termites in total, Fig 2Aii, 2C) and minor workers (14 trials, 420 termites in total, Fig 2Bii, 2C). This behavioural attraction means that in the experiments, their preference index PI (See methods) to *Fresh Colony soil* was closer to +1 (Fig 2C; p=0 for majors and p=0 for minors, Wilcoxon rank-sum test, Table S3). This attraction to soils was not a collective phenomenon, as individual termites showed similar preferences in the choice assay (Fig S1). Thus, major and minor workers were individually (Movie S3; Majors: 40 trials, 40 termites; minors: 40 trials, 40 termites) and collectively attracted to *fresh colony soil* against *control soil*, suggesting that *fresh colony soil* contained physical or chemical cues that attracted nestmates.

**Figure 2:**
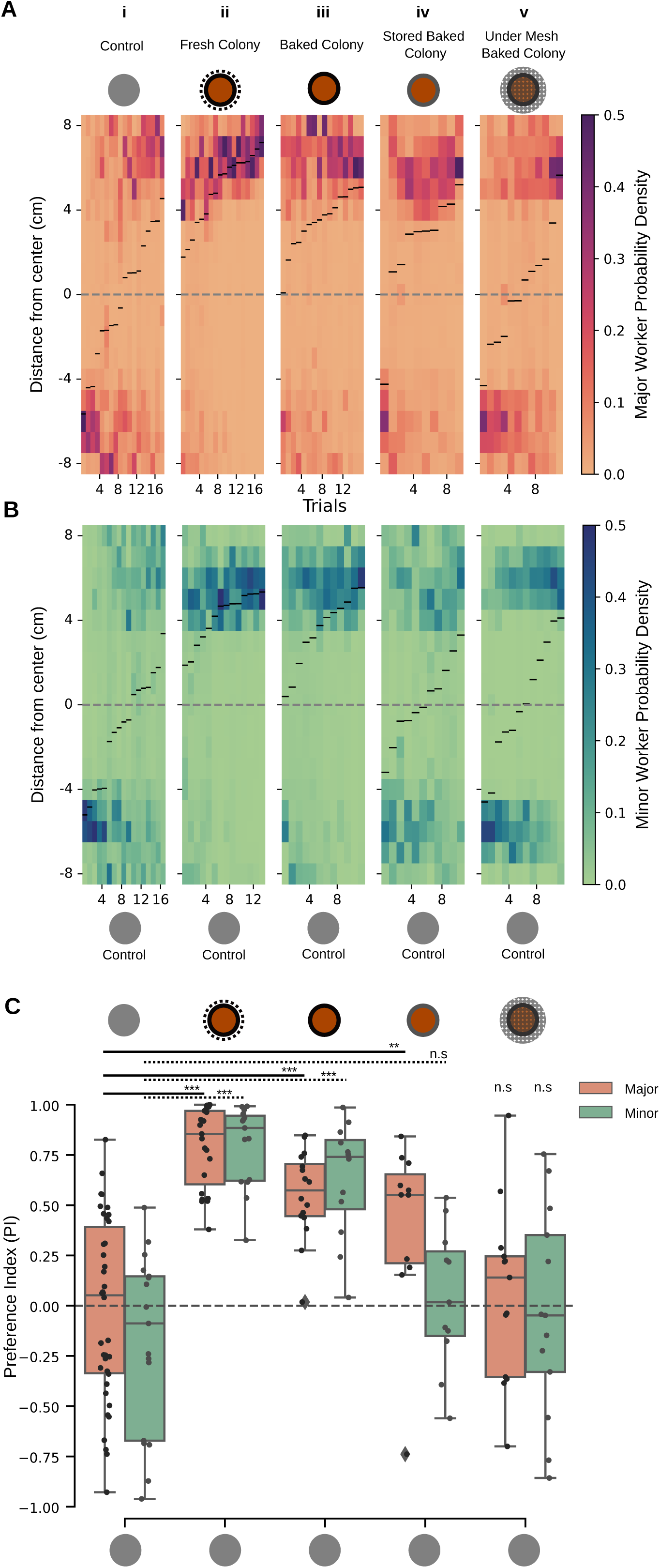
Soil at the site of repair attracts termites and contains volatile and non-volatile chemicals. Tapestry heatmaps show the spatial probability density of termites in the arena during the 50-55 min window across multiple trials. The color bar indicates the number of termite detections at each y-coordinate, and the black line in each strip denotes the median position for that trial. The y-axis is centered at 0 cm (release point) and extends ±8.5 cm. **A-B)** Major (A) and minor (B) worker responses across five conditions (left to right): i) Control vs Control; ii) Fresh Colony vs Control; iii) Baked Colony vs Control; iv) Stored Baked Colony vs Control; v) Baked Colony under mesh vs Control. **C)** Boxplots of the Preference Index (PI) for each trial over the same 5 min window. Majors are shown in orange, minors in green. Trial numbers may differ slightly from heatmaps, as all trials were included in the statistical analysis. Trial numbers (box plots): **i)** Majors: 39, Minors: 17; **ii)** Majors: 20, Minors: 14; **iii)** Majors: 17, Minors: 12; **iv)** Majors: 10, Minors: 11; **v)** Majors: 13, Minors: 13. Significance assessed using the Wilcoxon rank sum test; **p < 0.01, ***p < 0.001, *n.s.* = not significant (see Table S3 for details).

### Nestmates are attracted to both volatile and non-volatile pheromones in Fresh Colony Soil

We next tested whether attraction required airborne (volatile) signals or contact-dependent (non-volatile) cues in the freshly processed soil. In these experiments, the soil was baked at 60°C for 48-56 hours to eliminate volatile cues, and we measured the attraction of the termites either to the *baked colony soil* lacking volatile cues, or *fresh colony soil* containing both volatile and non-volatile cues.

### Baking process eliminates volatile cues leaving behind only non-volatile chemical cues

Major and minor workers were attracted to *baked colony soil* over *control soil* (Fig 2Aiii, 2C; Majors: 17 trials, 510 termites, p=0; Fig 2Biii, minors:12 trials, 360 termites, p=0 Wilcoxon rank-sum test, Table S3). Thus, even after elimination of volatile cues, persisting non-volatile cues in the baked colony soil were sufficient to attract the termites. Under natural circumstances, the termite mound is often subjected to prolonged heat under the sun. To test if the odor signals from the baked soil last over long durations, we compared the responses of termites to *Stored Baked Colony soil* (see Methods), which was stored under laboratory conditions for as many as nine months, against *control soil*. Over such time scales, only major workers demonstrated a clear preference for the *Stored Baked Colony soil*, whereas minor workers showed no distinct preference (Majors: 10 trials, 300 termites, p=0.004; Minors: 11 trials, 330 termites, p=0.335, Wilcoxon rank-sum test, Table S3) (Fig 2Aiv, 2Biv), which suggested that major termites are more sensitive to these cues over a long term. To determine the efficacy of the baking process in eliminating volatile compounds, the *Baked Colony soil* was placed under a mesh (see Methods, Fig 1D, E). Absence of any termite attraction towards this patch confirmed that volatile cues were effectively removed post-baking (Fig 2Av, 2Bv; Majors: 13 trials, 390 termites, p=0.82; minors:13 trials, 390 termites, p=0.287; Wilcoxon rank-sum test, Table S3). Together, these tests showed that the baking process eliminates volatile cues but maintains attraction towards non-volatile cues over the entire timeframe of our experiments, and possibly even longer.

### Fresh colony soil contains volatile chemical cues

When presented with a choice between *Fresh vs Baked Colony soil*, termites were attracted to the fresh colony soil (Fig 3Bi, 3Bi; Majors: 20 trials, 600 termites, p=0; minors:16 trials, 480 termites, p=0; Wilcoxon rank-sum test), consistent with the hypothesis that the *Fresh Colony soil* contains attractive volatile compounds, in addition to the non-volatile cues.

**Figure 3:**
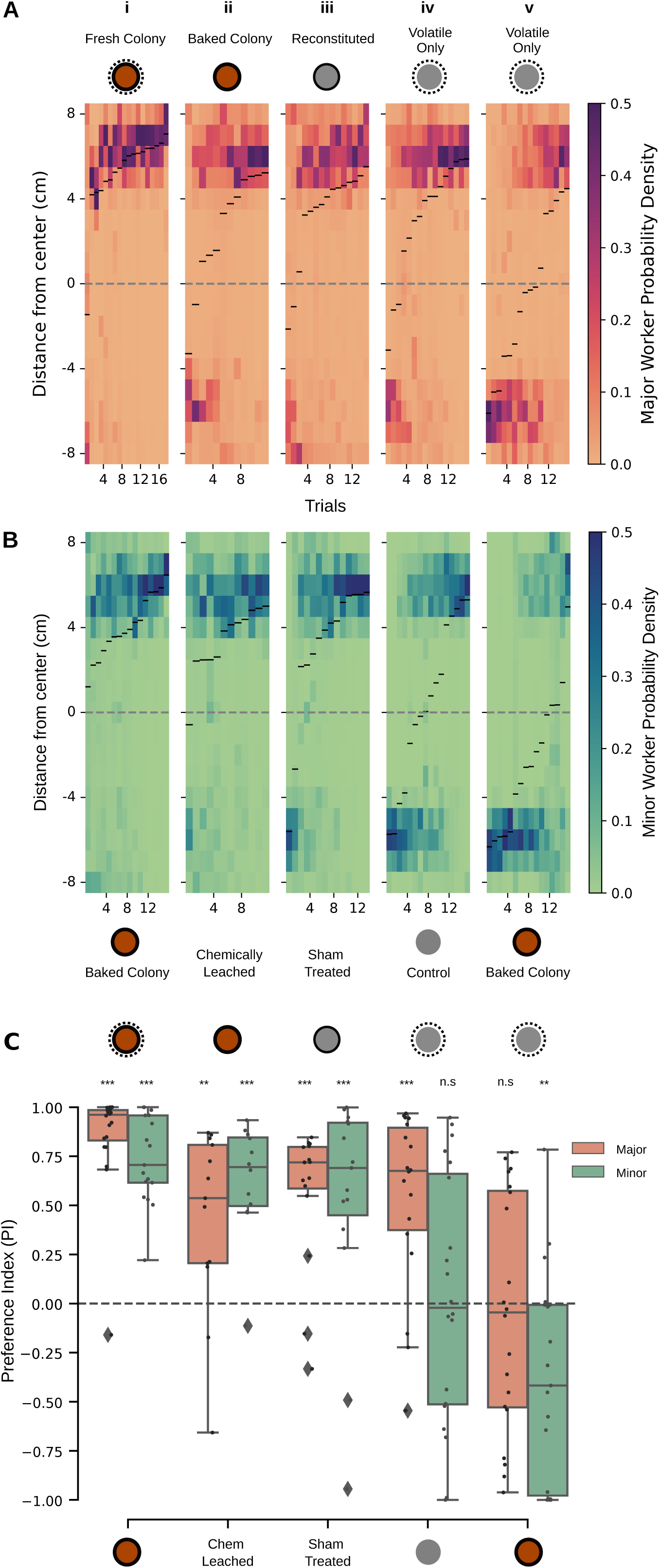
The chemical extract from Fresh Colony Soil is sufficient to impart attraction to Control soil. Tapestry heatmaps show termite spatial probability distributions during the 50-55 min window, aggregated over multiple trials. The color bar indicates the number of termite detections at each y-coordinate; median positions are marked by black lines. **A-B)** Representative tapestry heatmaps showing the probability density of termites in the arena during the 50–55 min time window. The color bar indicates the number of termite detections at each coordinate across trials. The black line on each trial is the median point for the trial. The y-axis is centred at 0 cm (release point) and extends to ±8.5 cm. From left to right, columns represent different experimental conditions: **i)** Fresh Colony Vs Baked Colony soil (Majors: 20 trials; minors: 16 trials), **ii)** Baked Colony soil Vs Chemically-leached colony soil (Majors: 13 trials; minors: 11 trials), **iii)** Reconstituted Vs Sham treated soil (Majors: 15 trials; minors: 14 trials), **iv)** Fresh Colony under mesh (only volatile cues) Vs Control soil (Majors: 18 trials; minors: 20 trials), **v)** Fresh Colony under mesh (Volatile only) Vs Baked Colony soil (Majors: 20 trials; minors: 19 trials). **C)** The PI for each trial is plotted as a single point for the 5 min window as Boxplots for Majors(orange), minors(green). Wilcoxon rank sum test Table S3, when compared against the control experiments. Statistics: **p<0.01, ***p<0.001, n.s-not significant compared again Control Vs Control distributions, Wilcoxon rank sum test (See Table S3).

### Chemical cues are sufficient to elicit attraction

#### Influence of Non-Volatile cues

To ascertain the chemical nature of the cues, we solvent-extracted non-volatile compounds from termite-processed soil (see Methods). In the choice assay between *Baked colony soil* and *chemically-leached colony soil*, termites were not attracted to the chemically-leached soil. Thus, the extraction process diminished the pheromone signal in the *chemically-leached colony soil* relative to *Baked colony soil* (Fig 3Aii, 3Bii; Majors: 13 trials, 390 termites, p=0.003; minors:11 trials, 330 termites, p=0; Wilcoxon rank-sum test). Next, we added the solvent extract to *unprocessed soil* to test if it acquired the ability to attract nestmates, compared to a *sham-treated control* with solvent alone. In these experiments, termites were attracted to the non-volatile *reconstituted soil* rather than the *sham-treated soil* (Fig 3Aiii, 3Biii; Majors: 15 trials, 450 termites, p=0; minors:14 trials, 420 termites, p=0.0004; Wilcoxon rank-sum test, Table S3). These experiments confirmed that termites add chemical pheromones to the soil, and the chemical extract of these cues was sufficient to generate attraction towards soil that was not processed by termites.

#### Influence of Volatile Chemicals

Next, in the *mesh choice assay* (See methods: Mesh Choice Assay, Fig 1D, E), a porous metal mesh separated the termites from the soil placed beneath allowing only volatile cues to pass through. The termites then had to choose between the volatile cues from *Fresh colony soil (Volatile Only soil)* and the unprocessed *Control soil* (Movie S4). On both patches, *Control soil* was placed on the top of the mesh to ensure that termites had soil to work with (See methods). Under these conditions, the major workers were attracted to the *Volatile Only soil* (Fig 3Aiv; Majors: 18 trials, 540 termites, p=0 Wilcoxon rank-sum test); thus, volatile cues alone were attractive for these termites. In contrast, however, minor workers were equally distributed at both ends (Fig 3Biv; Minors: 20 trials, 600 termites, p=0.247; Wilcoxon rank-sum test, Table S3). Thus, volatile cues from termite processed soil were attractive only for the major workers, but not the minor workers. This suggests that major workers are likely recruited by their nestmates using volatile cues deposited in the soil.

#### Major and minor castes exhibit different responses

Do major and minor termites respond differently to the combination of volatile and non-volatile cues present in the processed soil? To address this question, we presented termites with a choice between volatile and non-volatile cues. At one end of the arena, we placed *Volatile Only soil*. At the other end, we placed *unprocessed soil* under the mesh but overlaid it with *Baked colony soil*, with which they could contact. This ensured that at one end, the termites were presented with volatile cues but no contact (non-volatile) cues, whereas at the other end, they had access to contact cues but no volatile cues. The presence of control soil at both ends also ensured comparable levels of humidity. In this experiment, major termites were distributed equally at both ends (Fig 3Av; Majors: 20 trials, 600 termites, p=0.8, Wilcoxon rank-sum test), whereas minor termites only aggregated at the end containing non-volatile cues (Fig 3Bv; Minors: 16 trials, 480 termites, p=0.004; Wilcoxon rank-sum test).

Together, these experiments showed that major workers were attracted to volatile cues in isolation, while minor workers were not. However, when a choice between non-volatile cues and volatile ones was given, minor workers were attracted to the non-volatile cues, and major workers showed no preference, indicating caste-specific responses to soil-borne cues.

## Discussion

Unlike most social insects, workers and soldier termites lack image-forming eyes and rely heavily on pheromone cues that they deposit on the structure or emit in the air (22–24). To identify the pheromones and their active components, we need experimental assays capable of accurately measuring the behavioural changes in response to various chemical environments. The *choice assay* described in this paper can measure the subtle behavioural effects of different components within a pheromone complex. Experiments conducted with the choice assay and its variations yielded multiple findings. First, the soil processed by termites contains non-volatile and volatile chemicals which attract their nestmates, promoting aggregation and construction in that soil (Fig 2Ai-iii, Bi-iii). *Unprocessed control soil* became attractive to major and minor workers when infused with chemical extracts from *fresh colony soil* (Fig 3A ii-iii, B ii-iii). This demonstrated that termites deposit chemical cues when they process the soil. Second, workers were more attracted to *fresh colony soil* containing both volatile and non-volatile cues, as compared to the *baked colony soil* which lacked volatile cues (Fig 3Ai, Bi). Thus, volatile cues enhance the attraction of the soil for both major and minor workers, over and above non-volatile cues contained in the *baked colony soil*. Third, in a modified choice assay, *fresh colony soil* was enclosed in mesh to allow volatile cues to pass through while preventing contact with non-volatile cues. In these experiments, both volatile and non-volatile cues individually attracted major workers (Fig 3Aii, iv), but minor workers were attracted to non-volatile cues alone (Fig 3Bii,iv). Fourth, major and minor workers showed caste-specific responses to volatile and non-volatile cues. Whereas major workers were attracted to purely volatile cues regardless of the presence of non-volatile cues, the minor workers were attracted to volatile cues only when non-volatile cues were present. Indeed, to be attractive to minor workers, the volatile cue needed to be colocalized with the non-volatile cue.

In the presence of these cues, termites stayed and built in the soil processed by their nestmates, which strongly indicates they constitute a ‘clay’ (or ‘cement’ or ‘construction’) pheromone which has been the subject of some debate (25–28). Because major and minor workers respond differently to volatile and non-volatile cues, these chemicals help guide their specific roles in building and repair. Consistent with the Grassé’s *stigmergy hypothesis*(*14*), this communication does not occur directly between individual termites, but is mediated by the soil in which the cues are deposited (29).

Under natural circumstances, major and minor workers of mound-building termites are thought to serve different functions. For instance, in *Macrotermes bellicosus* (30), major termites forage widely and are involved in diverse tasks, including construction and maintenance, and minor termites are mostly nest-bound, involved in emergency construction and tending the queen. In *Odontotermes obesus,* both major and minor workers collectively participate in mound construction and maintenance activities (31, 32). Their specific roles in this process under natural conditions are hard to decipher because the mounds are closed structures, and termites resist any attempt to film within the mound by depositing a layer of soil on foreign objects. Nevertheless, the response of major and minor workers to volatile vs non-volatile cues is different but complements each other. Because major workers are attracted to volatile cues, they are likely recruited from a distance. However, minor workers are attracted to volatile cues only if spatially co-localized with non-volatile cues. In the context of mound repair, this suggests that the first responders to a breach in the mound are likely to be minor workers, who sense the breach using either light, acoustic signals or air disturbance from the site of the breach (32). Their arrival at the site and subsequent repair work cause volatile and non-volatile cues to accumulate, recruiting major workers from a distance.

To test this prediction, we reanalysed existing cone assay data on termite repair behaviour (31). This revealed that the initial response (in the first 40 minutes) was dominated by minor workers (Fig. S2), consistent with our findings and earlier observations in *M. bellicosus* (33) in which worker ratios evolved during foraging and minor workers were the initial builders (30). To test if major workers are involved in long-term repair, we offered mound soil left exposed for over a month (See SI Methods) and observed that major workers were still attracted to this soil (Fig S3), and perhaps involved in mound maintenance.

Together, these experiments provide insights into the nuanced communication system of termites in their natural context. The methods outlined here can be extended to identify the specific pheromones which enable termites to stigmergically construct or repair mounds. These experiments may also benefit roboticists aiming to develop platforms for collective construction (34) as biorobotics moves beyond simple swarm algorithms toward more sophisticated task allocation systems. Our results suggest three key principles for such systems. First, functional specialization via differential sensory tuning among otherwise identical agents. Second, chemical gradients as spatial organisers for sequential tasks, and third, long-lasting non-volatile cues that promote site fidelity across multiple agent cohorts. These ideas could inform the design of decentralized, construction-capable robotic swarms.

## Materials and Methods

### Termite collection

*Odontotermes obesus* termites were collected from 10 mounds located in the National Centre for Biological Sciences (NCBS) and Gandhi Krishi Vignana Kendra (GKVK) campuses in Bengaluru, India. To simulate damage, a ∼4 cm hole was made in each mound and covered with a 50 mL centrifuge tube. Termites entered the tube and lined its interior with soil. After more than half the tube was coated (typically in 1.5-6 hours), it was sealed. The trapped termites were used in experiments, and the deposited soil was used as *fresh colony soil*. To reduce inter-caste variability, major and minor workers (Fig 1 B) were separated and tested independently.

### Moisture measurement

Soil moisture was measured using the gravimetric method: wet soil was weighed, dried at 60 °C for 48-56 hours, and weighed again. Moisture content was calculated as the ratio of water mass (wet – dry weight) to dry weight, yielding a w/w percentage (gram/gram) of water content.

### Soil treatments

To compare termite responses to different chemical cues in soil, we prepared a range of soil treatments (Table S1). All soils except *Fresh Colony soil* were oven-dried and rehydrated to 25% w/w moisture to ensure consistency across treatments. *Fresh Colony soil* was tested at its natural moisture level (see SI Methods, Fig. S4).

**Control soil** was collected from the mound’s vicinity and not processed by termites. **Fresh Colony soil** was directly collected from the mound, and contained both volatile and non-volatile termite-derived compounds and was not dried.

**Baked Colony soil** was Fresh Colony soil that had been oven-dried to remove volatiles, with moisture restored to 25% w/w.

**Stored Baked Colony soil** was Baked colony soil stored for up to 9 months to test for cue longevity.

**Chemically-leached Colony soil** was Fresh Colony soil subjected to solvent washes to remove chemical cues.

**Reconstituted soil** was Control soil infused with the extract from chemically leached soil, retaining non-volatiles but lacking physical termite processing.

**Volatile-only soil** was prepared by placing Fresh Colony soil below a mesh, allowing exposure to airborne cues only.

**Sham-treated soil** was Control soil exposed to the same chemical treatment as Reconstituted soil, without added cues.

**Mound soil** was naturally aged, termite-processed mound soil, not freshly collected or heat-treated.

Among these, only Control, Reconstituted, and Sham-treated soils were not processed by termites. All others had undergone direct interaction with the termites.

(See SI Methods for full preparation protocols and solvent extraction details.)

**Figure 4:**
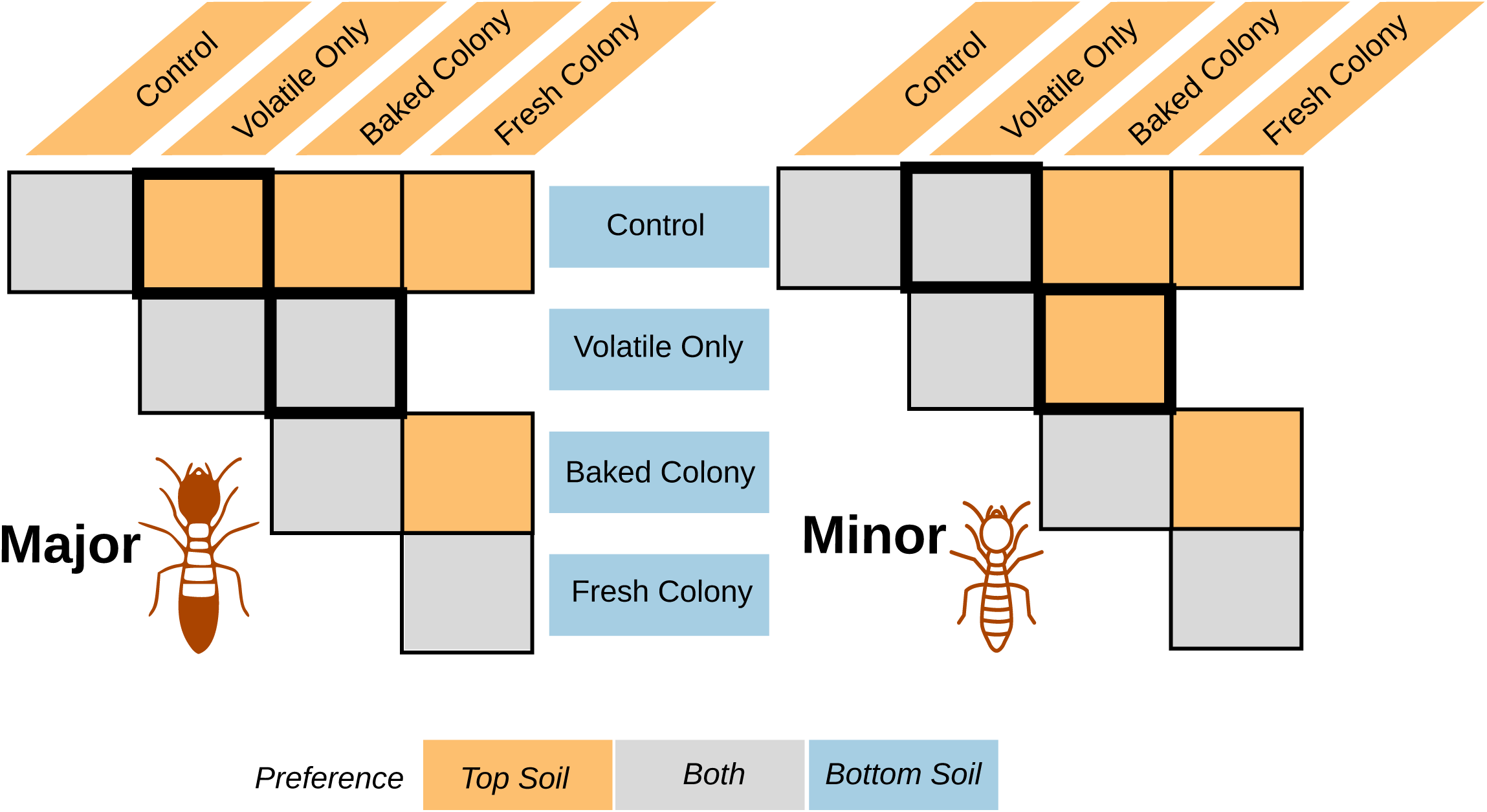
Caste-specific differences in soil preferences across treatments. Matrix summarizing termite preferences across experiments for major and minor workers of *Odontotermes obesus*. Each cell indicates the dominant caste-specific preference in a given assay: preference for the top soil (orange), bottom soil (blue), or no significant preference (gray). Columns and rows represent different experimental pairings. Black outlines highlight conditions where major and minor worker preferences diverged.

### Experimental Setup

#### The Choice assay

Near each end of a rectangular plexiglas arena (170 mm x 70 mm x 80 mm; Fig 1C), we placed two circular patches of soil (30 mm diameter, 1.5 mm thickness) 80 mm apart. The starting moisture content of each soil patch was 25% w/w of dry soil (Fig S3). Unless mentioned otherwise, 30 termites of either major workers or minor workers were introduced at the center of the arena and allowed to freely move between the two soil patches. The arena was illuminated using two diffuse LED light sources on each side, cumulatively providing a light intensity ∼1000 lux. The collective movement of termites is filmed from an overhead view using a DSLR camera (Nikon D5200, D7500, 1080p) at 30 frames per second. Experiments were conducted in a room where the temperature was 25 *±* 2 °C and 65 *±* 10% RH. Our preliminary observations showed that steady-state distributions were reached at 40 minutes after the termites were introduced in the arena for the control vs control experiments. Based on this observation, we set the total time of filming to 60 minutes and data in the 50-55 minute period was quantified as a steady-state choice.

#### Mesh Choice assay

To test termite responses to volatile cues alone, we constructed a two-layered arena with the same dimensions as the standard choice assay (Fig. 1D). The floor of the top panel included a mesh (300 μm grid) stretched across a 30 mm diameter depression. This mesh prevented physical contact with soil beneath while allowing volatiles to diffuse upward. Test soils were placed below the mesh, and a semi-circular patch of soil was positioned above. Termites were introduced on the top panel. Depending on the test, different experimental or control soils were used. Aggregation or building on the mesh surface was interpreted as a response to airborne cues. (See SI Methods for full construction details and odor containment controls.)

#### Data analysis

A 5-minute steady-state window (50-55 min) was digitized from each 60-minute video using *Trex* (*35*). Termite positions were tracked along the arena’s y-axis, and density heat maps were averaged over 300 seconds to generate a composite spatial distribution. Median termite positions were marked with black lines (Fig. 1F), and trials were plotted side-by-side in a tapestry format, sorted by ascending medians for visual clarity.

All code was written in Python, with visualizations generated using Seaborn and Plotly Express. (See SI Methods for axis definitions and processing steps.)

#### Individual termite experiments

We analyzed the full 0-30 minute duration, calculating the we used a tapestry plot where the x-axis represented the number of trials and the y-axis represented the superimposed average position over the entire trial, from 0 to 30 minutes. As with other plots, the median position was marked in black, and the trials were sorted and arranged in ascending order of the medians (Fig S1). (See SI Methods for trial criteria and exclusion rationale.)

#### Preference Index

The arena was divided into two and the preference index was calculated as the ratio of the difference of termites on the top and bottom sides of the well to the sum of termites on the top and bottom.

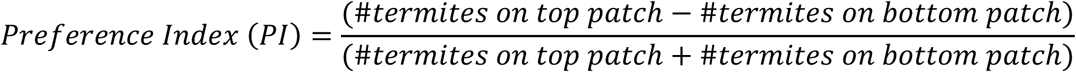

The value ranges between -1 and +1. If the PI value is positive, then the top Patch has a preference over the bottom Patch. Values closer to 1 and -1 denote a strong preference and closer to zero denote no preference.

### Statistics

The normality of Preference Index (PI) values was assessed using the *Shapiro-Wilk* test. As some distributions were non-normal, we used the non-parametric *Wilcoxon rank-sum* test for comparisons between independent groups, and the *Wilcoxon signed-rank* test to evaluate patch preference within trials. Unless specified, each experimental set was tested against a control where both patches contained control soil (see Figs. 2C, 3C; Tables S2-S3). Levels of significance are denoted as *\* p < 0.05, **p < 0.01, and ***p < 0.001*. Full statistical results, including test statistics and exact *p*-values for all treatment caste combinations, are reported in SI Tables S2 and S3.

## Data Availability

Data analysis and recreation of figures and codes are available on request

## Author contributions

S.S.R. designed and performed experiments, analyzed data, and wrote the manuscript. S.P.S. provided guidance, supervised the research, and wrote the manuscript.

## Acknowledgements

We thank Pavan Kumar Kaushik and members of Sane lab for discussions, and the Mechanical, Electrical and Electronics workshops, Civil department and gardening staff. S.S.R was supported by the Department of Biotechnology, Government of India (DBT/2018/NCBS/1158). S.P.S was supported by the Air Force Office of Scientific Research grants (AFOSR; FA2386-11-1-4057 and FA9550-16-1-0155) and the National Centre for Biological Sciences, Tata Institute of Fundamental Research, Department of Atomic Energy, Government of India. This project was originally funded by the Human Frontier Science Program (RGP0066/2012) for a project in which S.P.S was a co-investigator.

## Supplementary Information

**Fig S1:**
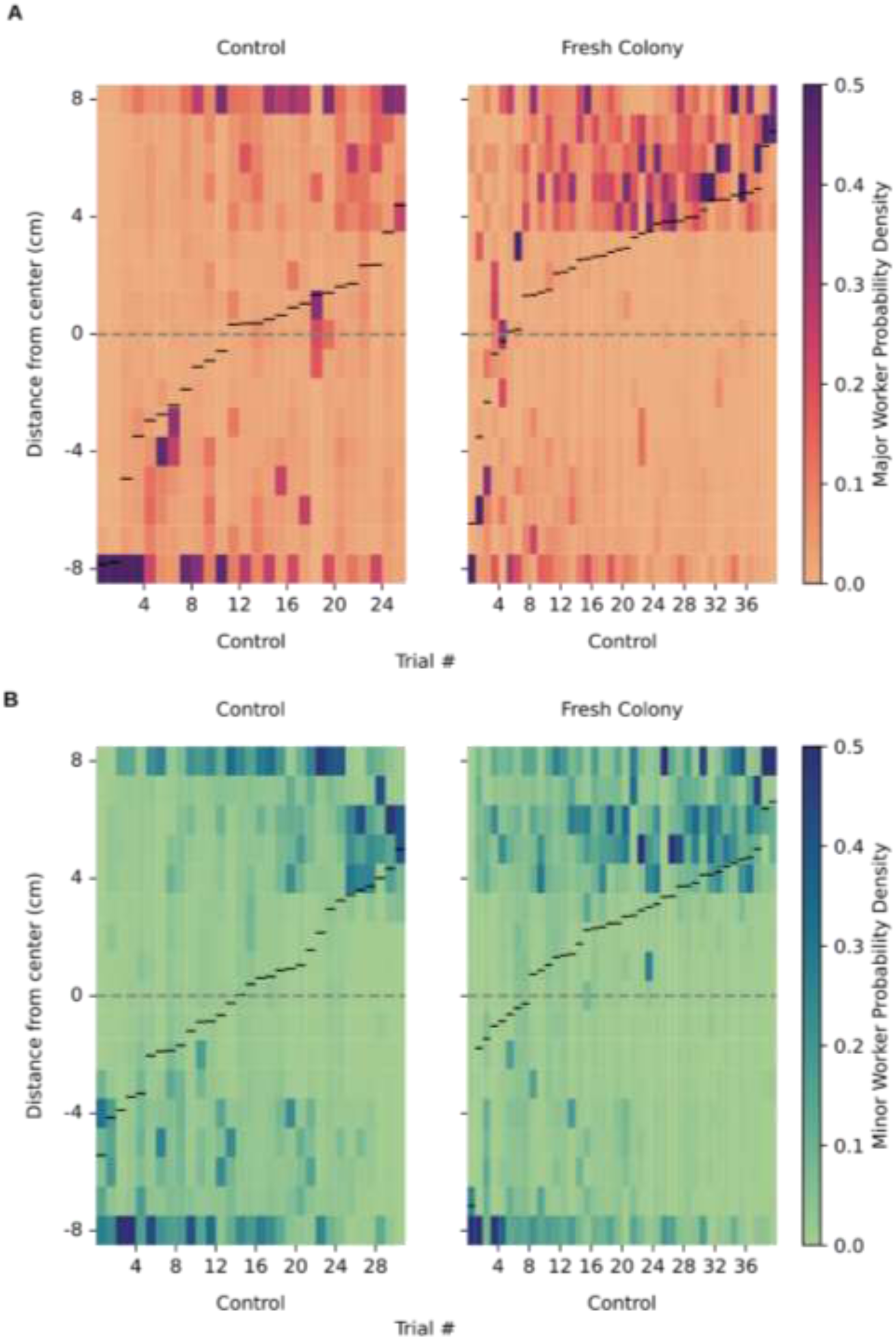
Individual termites prefer the fresh colony soil patch.: Tapestry heatmaps showing individual termite trajectories in a choice assay between fresh colony soil and control soil (Movie S3). Each heatmap represents the collapsed average position of a termite over the full 30-minute trial. Unlike group-level experiments that showed steady-state patterns, individual termite movements were not split into time segments because their behavior never reached a steady state. (A) **Major workers:** 26 trials (control vs. control), 40 trials (fresh colony vs. control). (B) **Minor workers:** 31 trials (control vs. control), 40 trials (fresh colony vs. control). These results suggest that individual termites, like groups, exhibit a consistent preference for the fresh colony soil.

**Fig S2:**
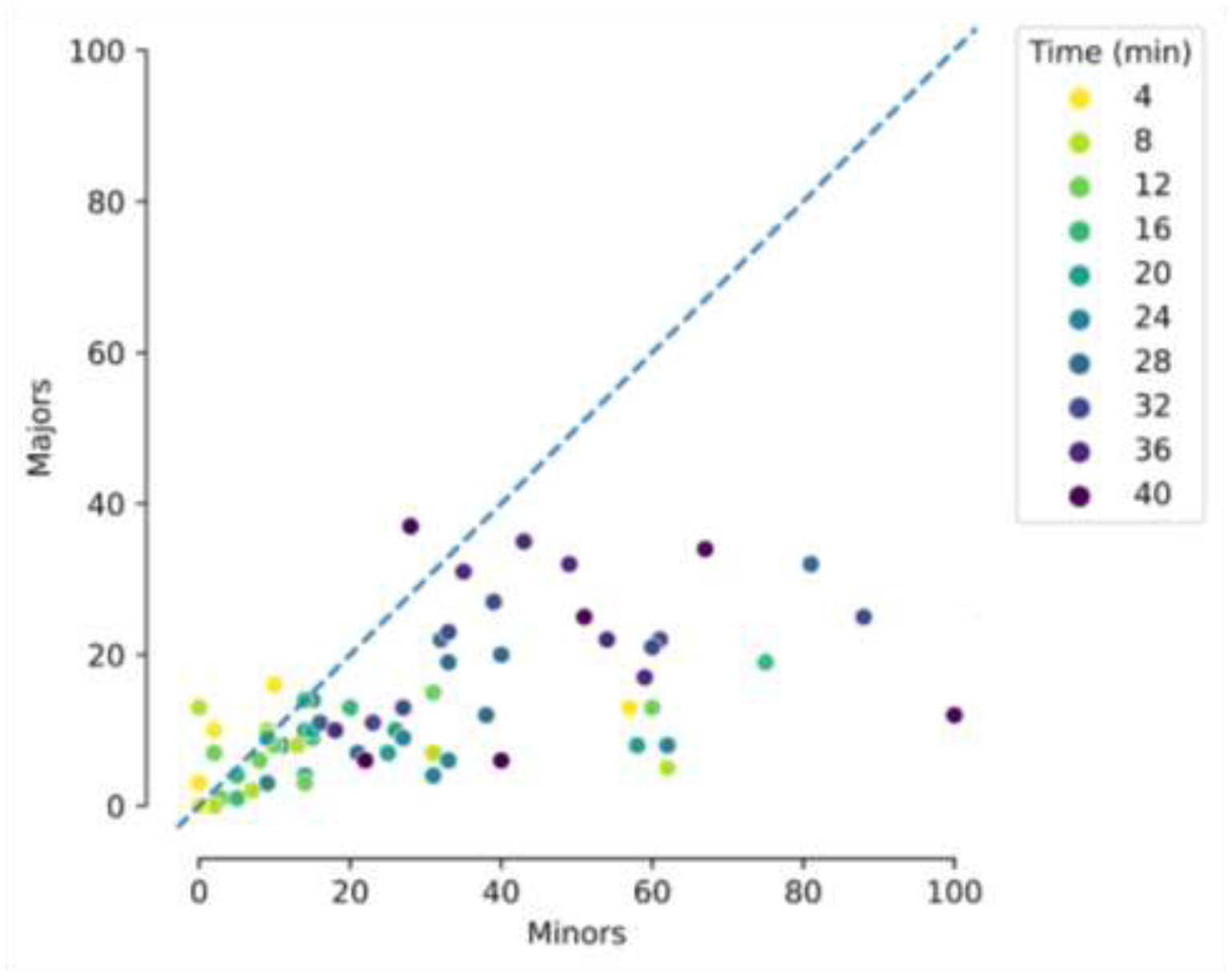
Caste composition dynamics during mound repair. During mound repair, the relative abundance of worker castes shifts over time. Early repair phases tend to be dominated by minor workers, whereas later phases feature an increasing proportion of major workers. Each point represents a time bin within a trial (n = 7 trials). The x-axis shows the absolute number of minor workers; the y-axis shows the absolute number of major workers. Colors represent different time points across trials. The diagonal line (y = x) indicates equal representation of majors and minors. As time progresses, data points tend to shift above the diagonal, indicating a major-biased composition (31).

**Fig S3:**
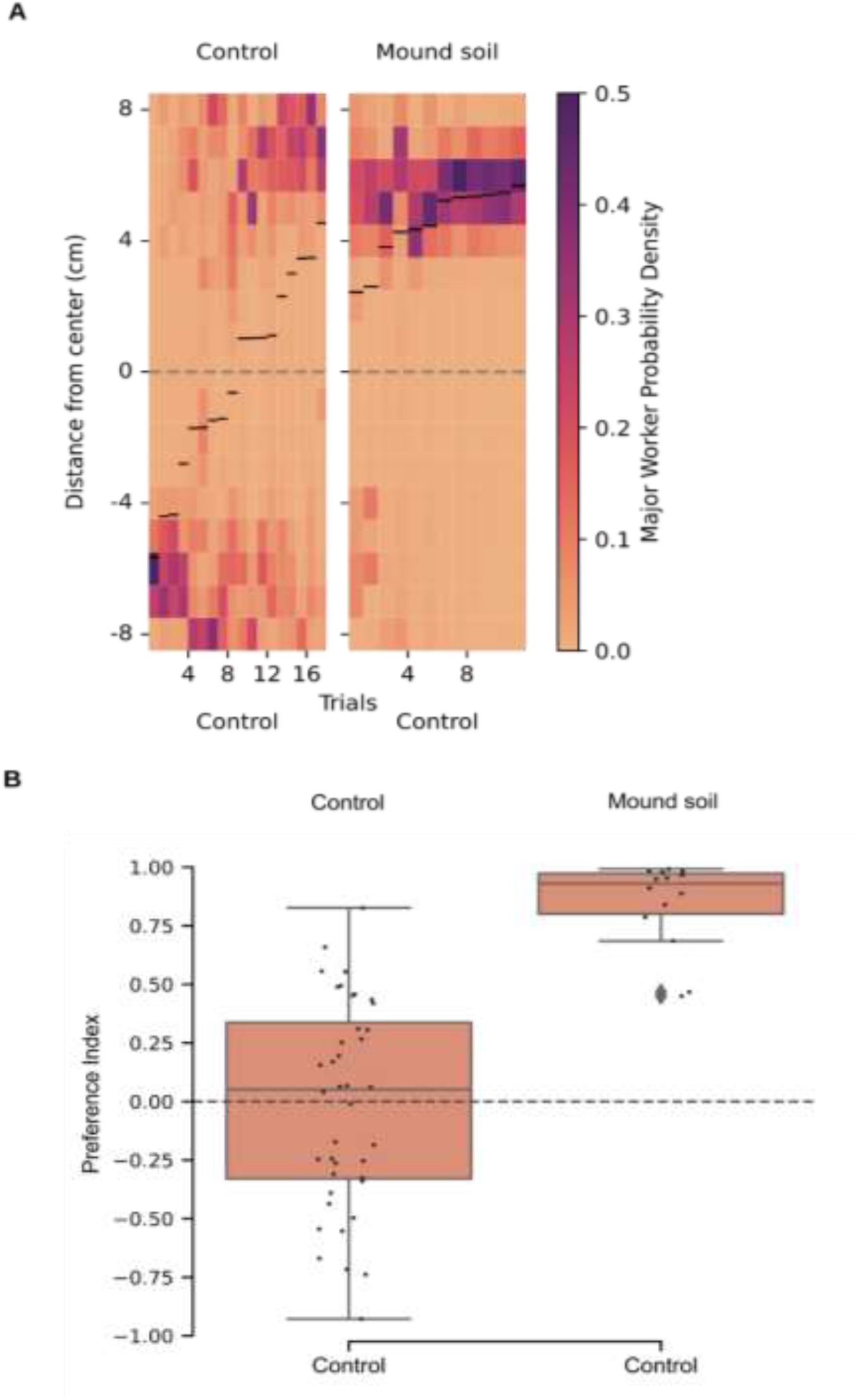
Mound soil remains attractive to major workers after sun exposure. **A)** Even after one month of sun exposure, mound soil remains attractive to major workers when compared to control soil (see SI Methods for soil preparation details). Tapestry heatmaps of Control Vs Control and mound soil Vs control soil. **B)** Boxplots show the Preference Index (PI) for two conditions: control vs. control (n = 39 trials, left) and mound soil vs. control (n = 12 trials, right). The mound soil continues to elicit a significant attraction (p=0, Wilcoxon rank sum test) response, suggesting persistent chemical cues despite environmental exposure.

**Fig S4:**
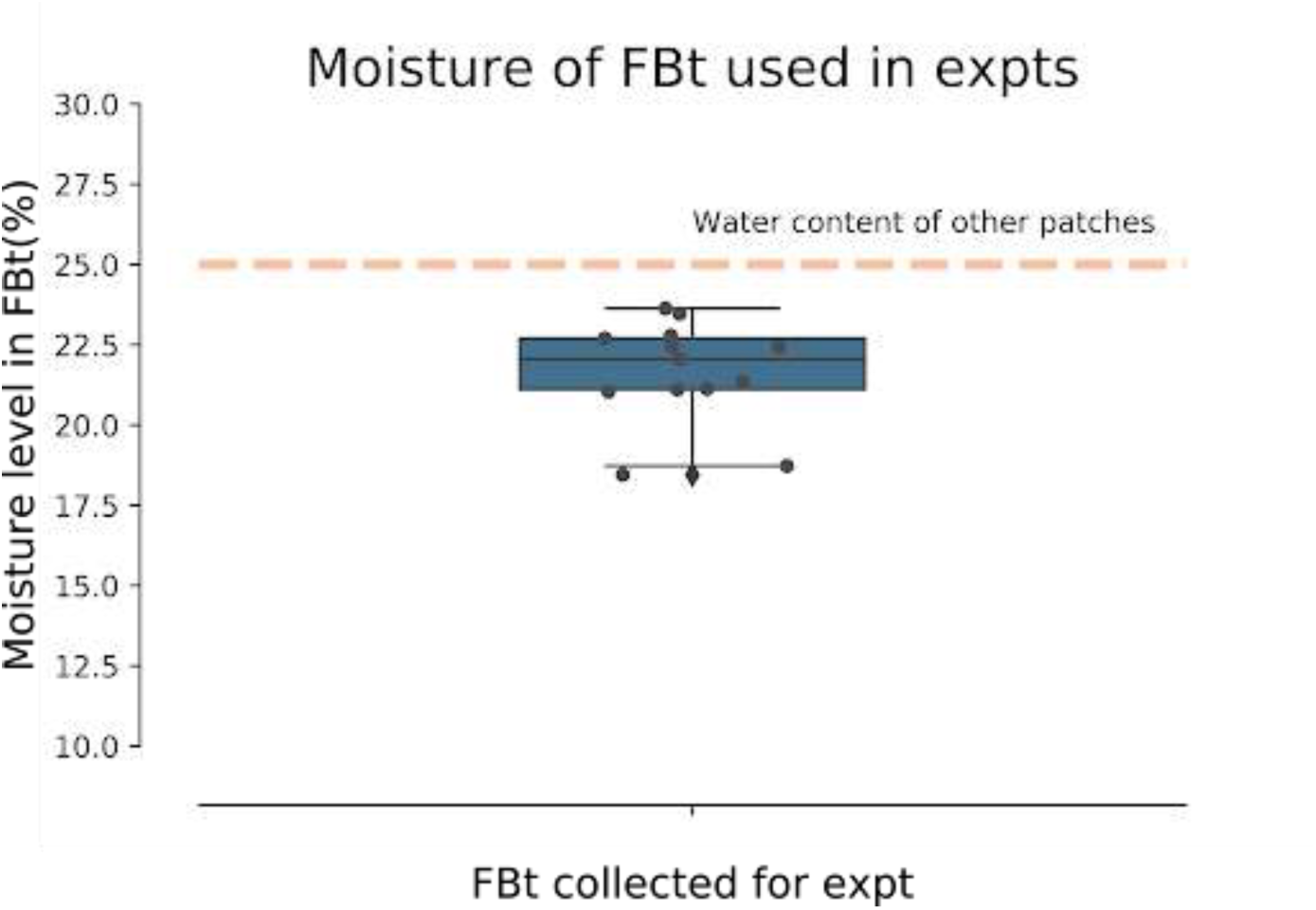
Post hoc moisture measurements of fresh colony soil: All soil patches were initially prepared at 25% moisture content (w/w of dry soil). Post hoc gravimetric analysis revealed that the fresh colony soil patch had a slightly lower moisture content, averaging 22% ± 2% (mean ± SD), likely due to external conditions or natural variability.

**Movie S1: Termites building at the site of repair.** Termites actively building at a breach in the mound. The video is shot and played at 25 frames per second.

**Movie S2: Choice assay with 30 termites:** Video of an experiment to measure termite movement within the choice assay. Left lane contains two patches of soil: *Fresh Colony soil* (top left patch, marked blue) and *Control soil* (bottom left patch). Both soil patches in the right well contain *Control soil* to test for side bias. Thirty major workers were introduced at the centre of each arena, and their choices tracked over 60 minutes. In the video displayed here, this 60 min duration is played in 45 seconds.

**Movie S3: Choice assay with individual termites.** Video of an experiment comparing *Fresh Colony soil* (top left patch, from a repair site marked orange) with *Control soil* (bottom left patch). As in Figure S2, the right well serves as a bias control, with both patches containing *Control soil*. A single major worker was introduced at the centre of each arena, and its choices were tracked over 30 minutes. In the video displayed here, this 30 min duration is played in 75 seconds.

**Movie S4: Mesh Choice assay to test for volatiles.** Video of an experiment presenting only volatile cues: *Fresh Colony soil under mesh* on one side (marked with a green sticker) and *Control soil* on the other. In each arena, 30 major workers were introduced at the centre and tracked for 60 minutes. The second well, with *Control soil* on both sides, serves as a bias control. In the video displayed here, this 60 min duration is played in 75 seconds.

## Supplementary methods

### Soil Handling and Moisture Control

All soil types were homogenised and shaped into uniform circular patches. Most soil patches were standardised to a moisture content of 25% w/w (relative to dry weight), except for the *Fresh Colony soil,* which was not rehydrated to preserve its natural volatile content. Post hoc measurements showed that its moisture content was 22% ± 2% (Fig. S4). These measurements helped assess variability in moisture content between termite-processed and unprocessed soils.

To avoid disturbing the termites during trials and to ensure that behavioural responses were not artefacts of moisture differences, we placed a duplicate, termite-inaccessible soil patch near each arena. These reference patches tracked moisture loss without influencing termite behaviour. Across all experiments, including Fresh Colony, Baked Colony, and Control soils, the average moisture loss during a trial was ∼4%, which was consistent across treatments.

### Soil Treatment Preparation

#### General Handling

Except for *Fresh Colony soil,* all soil samples were oven-dried at 60 °C for 48-56 hours, a temperature known to preserve organic matter while eliminating water and volatile compounds (37). This drying step enabled precise rehydration of all samples (except Fresh Colony) to 25% w/w during experiments. For Fresh Colony soil, natural moisture was preserved to maintain its volatile profile (Fig. S4).

#### Control Soil

Control soil was collected from ∼2 feet below the ground in the areas adjacent to termite mounds, ensuring no active termite or ant presence. The soil was broken down by pounding with a mallet and sieved through meshes of decreasing size. Only particles smaller than 2 mm were used, matching the texture of mound soil.

#### Fresh Colony Soil

*Fresh colony soil* was collected directly from the mound, particularly from freshly repaired sections, this soil is processed by termites and contained both volatile and non-volatile compounds. It was not subjected to oven-drying or any chemical treatment.

#### Baked Colony Soil

*Baked Colony Soil* was prepared by oven-drying *Fresh Colony soil* at 60 °C for 48-56 hours to remove volatiles. Prior to experimentation, it was rehydrated to 25% w/w water content.

#### Stored Baked Colony Soil

Portions of *Baked Colony soil* were stored in airtight containers for up to 9 months to evaluate the persistence of non-volatile cues over extended periods.

#### Chemically-Leached Colony Soil

*Fresh Colony soil* was treated with a sequence of solvent washes of increasing polarity: **hexane**, **acetone**, **acetonitrile**, **2-propanol**, and **methanol**. Each solvent was used to fully submerge the soil. Following this, the soil was rinsed three times with **distilled water,** then dried in a fume hood for 48 hours to reduce it from slurry to damp form. It was subsequently oven-dried at 60 °C for 48-56 hours to remove residual moisture. Supernatants from each solvent stage were collected and stored separately.

#### Reconstituted Soil

*Reconstituted Soil* is prepared by adding the supernatant from the *Chemically-leached soil* to *Control soil*, then processed identically (fume hood + oven-drying). The resulting substrate included non-volatile compounds from the mound but without termite processing.

#### Volatile-Only Soil

In this setup, *Fresh Colony soil* was placed beneath a mesh, and *Control soil* was placed above. The mesh prevented contact, allowing only airborne volatile cues to pass through. Termites could build on the top patch but had no access to the source of volatiles.

#### Sham-Treated Soil

*Sham-treated soil* was generated by subjecting Control soil to the same chemical treatment protocol as *Reconstituted soil* but without receiving any termite-derived compounds. This controlled for any attractive effects of the solvent process itself.

#### Mound Soil (Aged)

To assess the persistence of cues under natural conditions, a section of freshly built mound was monitored for one month before collection. This duration was chosen as longer observation periods were unreliable due to seasonal changes, mound growth variability, and risk of human interference. Additionally, as it was difficult to reliably collect minor workers from such sites, only major workers were used for these assays.

### Glossary of Soil Treatments

**Table S1:** Glossary of soil treatments used in the assays.

| Soil | Termite processed | Description |
| --- | --- | --- |
| Control | No | Unprocessed Soil |
| Fresh Colony | Yes | Mound Soil + Volatile + Non-volatile |
| Baked Colony | Yes | Mound Soil + Non-Volatile |
| Stored Baked Colony | Yes | Mound Soil + Non-volatile |
| Chemically-leached Colony | Yes | Treated (Mound Soil + Non-volatile) |
| Reconstituted | No | Treated (Unprocessed Soil + Non-volatile) |
| Volatile only | No | Unprocessed Soil + Volatile |
| Sham-treated | No | Treated (Unprocessed Soil) |

### Experimental Setup

#### Choice assay

The rectangular Plexiglas arena measured 170 mm × 70 mm × 80 mm (Fig. 1C). Two circular soil patches (30 mm diameter, 1.5 mm thickness) were placed 80 mm apart, approximately 20 times the body length of major workers (∼4 mm) and 26 times that of minor workers (∼3 mm) (Fig. 1B). The spacing ensured sufficient separation for detecting patch preference.

To ensure uniformity in patch size and shape, 3D-printed molds were used. Each mold was filled with the prepared soil mixture, inverted into the arena, and then removed to leave a clean circular patch in place.

#### Mesh Choice assay

We constructed a second arena with the same dimensions as the choice assay. The floor of this arena included a circular depression, 30 mm in diameter and a depth of 2 mm. This depression was covered with a mesh featuring a 300-micron grid (Fig. 1D). The mesh size was designed to prevent the termite legs and antennae from making physical contact with the soil beneath, ensuring that they were exposed only to volatile cues but unable to detect any non-volatile cues.

On the bottom panel (see Fig. 1D), the test soil was placed inside petri plates (35 mm diameter, 15 mm depth), positioned approximately 10 mm below the mesh. To ensure consistent placement and minimize lateral diffusion of odors, grooves matching the petri plate locations on the top panel were carved into a layer of packaging foam. This setup ensured that the petri plates remained in fixed positions throughout the experiment. The vertical columns formed by the foam and plates restricted odor diffusion to the vertical axis, allowing odors to rise upward into the top panel while preventing them from spreading across the bottom panel.

A semi-circular soil patch, 30 mm in diameter, was then positioned on top of the mesh (the top panel). The termites are introduced in the top panel. The soils in the bottom panel cannot be directly accessed by the termites. This setup allowed volatile compounds from the test soil to pass through the mesh. If termites were attracted by these volatile cues, they could aggregate on the soil patch placed above the mesh on the top panel. Based on the specific test conditions, either unprocessed (control) soil patches or other experimental soils were placed on the mesh surface, so termites could build on it. Unprocessed soil was placed beneath the mesh on the opposite side to control for moisture-driven biases in termite behaviour.

### Data Analysis

#### Data Inclusion and Recording

Trials showing no building activity were excluded from analysis. These non-building trials constituted less than 10% of the dataset and typically occurred in conditions where both patches were unattractive. Each experiment was recorded for 60 minutes. Due to camera memory limitations, this was done in three successive 20-minute video segments.

#### Digitisation and Coordinate Mapping

Videos were recorded at 30 frames per second (fps), providing high-resolution positional data for the observed termites. A 5-minute steady-state window between the 50th and 55th minute was selected for analysis. This window was chosen because termites exhibited stable behaviour by this time, and movement between patches had substantially reduced. We defined steady-state as the period when termite movement between soil patches decreased substantially, and visible building activity was initiated at one or both patches. Across trials, this phase was typically observed after 40 minutes, even in treatments where no attractants were present. This consistent onset of reduced movement and construction behaviour justified the use of the 50–55 minute interval as the representative window for final analysis and comparison across experimental groups. The final window was then digitised using *TRex* software (35), which extracted termite trajectories in each frame.

The y-axis of the arena (17 cm in length) was linearly mapped from –8.5 to +8.5, with 0 representing the centre, where termites were introduced. This consistent mapping allowed us to compare spatial distributions across trials. For each frame within the 5-minute sequence (300 seconds), a density heat map was generated along the y-axis. These were then averaged to produce a single composite probability density heat map per trial.

#### Trial Level Visualisation

To visualise termite distributions across all trials, the composite heat maps were arranged side-by-side in a columnar “tapestry” plot. Each column represented one trial, with the x-axis denoting trial number and the y-axis representing the mapped arena position. Trials were sorted in ascending order of their median termite position to facilitate interpretation. The median position across 30 termites for each trial was overlaid as a black horizontal line (Fig. 1F).

#### Individual termite experiments

Conducting experiments with individual termites proved especially difficult because in approximately 50% of the trials, termites either died or flipped onto their backs, becoming immobile. Such trials were excluded from the analysis. For the termites that were not excluded, the full 40-minute filming duration could often not be captured, either because some termites did not survive the entire 40 minutes, or because termites sometimes spent intermittent periods on their backs. Due to these limitations, all trials were analysed up to the 30-minute mark. The mean position along the y-axis was plotted and superimposed over the entire time-period. Because a preference index is not meaningful in this context, statistical analysis was conducted on the mean y-position for each termite. For individual termite experiments, we used a tapestry plot where the x-axis represented the number of trials and the y-axis represented the superimposed average position over the entire trial, from 0 to 30 minutes. As with other plots, the median position was marked in black, and the trials were sorted and arranged in ascending order of the medians.

### Statistics tables

Normality of Preference index values was assessed using the Shapiro-Wilk test (See Statistics table), which indicated that some groups were normally distributed while others were not. To compare the Preference Index (PI) values between independent treatment groups, we used the Wilcoxon rank-sum test (Mann-Whitney U test), a non-parametric test that considers both the magnitude and direction (positive or negative) of the continuous Preference Index (PI). For each pair of treatments, the test evaluates whether the two groups come from the same distribution by ranking all observations and calculating the sum of the ranks in each group. A significant result indicates that the distribution of PI values differs between the two groups, reflecting a potential difference in preference for the patches under different treatments (Fig 2C, 3C, Statistics table). Mostly, the distribution is compared against the *Control* Vs *Control* distribution for the main figures (See statistics table for complete comparisons).

To assess the statistical significance of patch preference within each trial, we used the Wilcoxon signed-rank test to compare the Preference Index (PI) values against an expected median of 0, which would indicate equal preference for both patches. The PI values represent the relative preference between two patches, where a positive or negative value reflects a bias towards one patch. The Wilcoxon signed-rank test, being non-parametric, is suitable for analysing the deviation from this expected value, particularly in cases where the assumption of normality may not hold. A significant result would suggest a preference for one patch over the other within the same trial. (See Full statistics table with all treatments: Table S2 and Table S3). Levels of significance are denoted as * for p<0.05, **for p<0.01, and *** for p<0.001. Unless specified, each experimental set was juxtaposed with a control where both patches were of control soil.

**Table S2:** Shapiro-Wilk Test for Normality and Wilcoxon Signed Rank Test for Within-Treatment Comparisons: This table displays the results of statistical analyses assessing the normality and comparing within-treatment groups. The Shapiro-Wilk test (‘shapiro_stat’ and ‘shapiro_p_value’) was performed to test the assumption of normality for each dataset. The “normal_distribution” column indicates whether the data is normally distributed (‘TRUE’ for normal, ‘FALSE’ for non-normal). The Wilcoxon signed-rank test was conducted, with the corresponding test statistic and p-value reported (‘Wilcoxon-statistic’ and ‘p_value’ columns).

| Treatments | termite caste | shapiro_stat | shapiro_p_value | normal_distribution | Wilcoxon-statistic | p_value |
| --- | --- | --- | --- | --- | --- | --- |
| Control Vs Control | Major | 0.970 | 0.376 | TRUE | 373 | 0.820 |
| Control Vs Control | minor | 0.926 | 0.187 | TRUE | 50 | 0.225 |
| Fresh Colony Vs Control | Major | 0.851 | 0.006 | FALSE | 0 | 0.000 |
| Fresh Colony Vs Control | minor | 0.864 | 0.035 | FALSE | 0 | 0.000 |
| Baked Colony Vs Control | Major | 0.946 | 0.434 | TRUE | 0 | 0.000 |
| Baked Colony Vs Control | minor | 0.925 | 0.330 | TRUE | 0 | 0.000 |
| Stored Baked Colony Vs Control | Major | 0.763 | 0.005 | FALSE | 9 | 0.064 |
| Stored Baked Colony Vs Control | minor | 0.957 | 0.636 | TRUE | 57 | 0.890 |
| Volatile loss from Baking | Major | 0.966 | 0.863 | TRUE | 38 | 0.970 |
| Volatile loss from Baking | minor | 0.971 | 0.921 | TRUE | 37 | 0.910 |
| Fresh Colony Vs Baked Colony | Major | 0.557 | 0.000 | FALSE | 1 | 0.000 |
| Fresh Colony Vs Baked Colony | minor | 0.923 | 0.186 | TRUE | 0 | 0.000 |
| Baked Colony Vs Chemically-leached Colony | Major | 0.866 | 0.058 | TRUE | 8 | 0.012 |
| Baked Colony Vs Chemically-leached Colony | minor | 0.848 | 0.035 | FALSE | 1 | 0.001 |
| Reconstituted Vs Sham-treated | Major | 0.719 | 0.000 | FALSE | 4 | 0.000 |
| Reconstituted Vs Sham-treated | minor | 0.760 | 0.002 | FALSE | 14 | 0.013 |
| Volatile only Vs Control | Major | 0.872 | 0.029 | FALSE | 9 | 0.001 |
| Volatile only Vs Control | minor | 0.934 | 0.207 | TRUE | 87 | 0.768 |
| Volatile Vs Baked Colony | Major | 0.921 | 0.177 | TRUE | 60 | 0.706 |
| Volatile Vs Baked Colony | minor | 0.916 | 0.145 | TRUE | 18 | 0.008 |
| Mound Vs Control | Major | 0.783 | 0.006 | FALSE | 0 | 0.000 |

**Table S3:** Wilcoxon Rank Sums Test for Comparison of Distributions: This table presents the results of the Wilcoxon rank-sum test, used to compare distributions across various termite caste and treatment groups. The table shows the pairwise comparisons between different treatment distributions (Distribution 1 vs. Distribution 2), along with the termite castes compared and the associated test statistics (statistic column) and p-values (p_value column).

| **Distribution 1** | **Distribution 2** | **Termite castes compared** | **statistic** | **p_value** |
| --- | --- | --- | --- | --- |
| Control Vs Control | Control Vs Control | Major-minor | 1.114 | 0.265 |
| Fresh Colony Vs Control | Control Vs Control | Major-Major | 5.813 | 0.000 |
| Fresh Colony Vs Control | Control Vs Control | minor-minor | 4.684 | 0.000 |
| Fresh Colony Vs Control | Fresh Colony Vs Control | Major-minor | 0.490 | 0.624 |
| Baked Colony Vs Control | Control Vs Control | Major-Major | 4.281 | 0.000 |
| Baked Colony Vs Control | Control Vs Control | minor-minor | 3.985 | 0.000 |
| Baked Colony Vs Control | Baked Colony Vs Control | Major-minor | -1.114 | 0.265 |
| Fresh Colony Vs Baked Colony | Control Vs Control | Major-Major | 5.614 | 0.000 |
| Fresh Colony Vs Baked Colony | Control Vs Control | minor-minor | 4.791 | 0.000 |
| Fresh Colony Vs Baked Colony | Fresh Colony Vs Baked Colony | Major-minor | 2.294 | 0.022 |
| Baked Colony Vs Chemically-leached Colony | Control Vs Control | Major-Major | 2.887 | 0.004 |
| Baked Colony Vs Chemically-leached Colony | Control Vs Control | minor-minor | 4.030 | 0.000 |
| Reconstituted Vs Sham-treated | Control Vs Control | Major-Major | 4.085 | 0.000 |
| Reconstituted Vs Sham-treated | Control Vs Control | minor-minor | 3.533 | 0.000 |
| Volatile only Vs Control | Control Vs Control | Major-Major | 3.595 | 0.000 |
| Volatile only Vs Control | Control Vs Control | minor-minor | 1.157 | 0.247 |
| Volatile only Vs Control | Volatile only Vs Control | Major-minor | 2.053 | 0.040 |
| Volatile Vs Baked Colony | Control Vs Control | Major-Major | -0.241 | 0.810 |
| Volatile Vs Baked Colony | Control Vs Control | minor-minor | 2.846 | 0.004 |
| Volatile Vs Baked Colony | Volatile Vs Baked Colony | Major-minor | -2.224 | 0.026 |
| Volatile loss from Baking | Control Vs Control | Major-Major | 0.222 | 0.824 |
| Volatile loss from Baking | Control Vs Control | minor-minor | 1.063 | 0.288 |
| Volatile loss from Baking | Volatile loss from Baking | Major-minor | 0.000 | 1.000 |
| Mound Vs Control | Control Vs Control | Major-Major | 4.841 | 0.000 |
| Stored Baked Colony Vs Control | Control Vs Control | Major-Major | 2.878 | 0.004 |
| Stored Baked Colony Vs Control | Control Vs Control | minor-minor | 0.963 | 0.336 |
| Stored Baked Colony Vs Control | Stored Baked Colony Vs Control | Major-minor | 2.607 | 0.009 |

